# Measurements of acoustic radiation force of ultrahigh frequency ultrasonic transducers using model-based approach

**DOI:** 10.1101/2021.01.17.427015

**Authors:** Sangnam Kim, Sunho Moon, Sunghoon Rho, Sangpil Yoon

## Abstract

Even though ultrahigh frequency ultrasonic transducers over 60 MHz have been used for single cell level manipulation such as intracellular delivery, acoustic tweezers, and stimulation to investigate cell phenotype and cell mechanics, no techniques have been available to measure actual acoustic radiation force (ARF) applied to target cells. Therefore, we have developed an approach to measure ARF of ultrahigh frequency ultrasonic transducers using theoretical model of the dynamics of a solid sphere in a gelatin phantom. To estimate ARF at the focus of 130 MHz transducer, we matched measured maximum displacements of a solid sphere with theoretical calculations. We selected appropriate ranges of input voltages and pulse durations for single cell applications and estimated ARF were in the range of tens of pN to nN. FRET live cell imaging was demonstrated to visualize calcium transport between cells after a target single cell was stimulated by the developed ultrasonic transducer.

---

Single cell engineering has great potential to visualize active molecular events during cell cycle while studying cell-cell interaction and sort intact cells without labeling^1–7^. While laser has been used for single cell level manipulation due to its short wavelength, photobleaching and thermal damage to cells may affect cell viability and functional and phenotypical changes after the manipulation^8, 9^. As the fabrication technique of ultrahigh frequency ultrasonic transducers ranging from 60 MHz to 300 MHz has emerged last two decades^10–12^, the wavelength of ultrasound are comparable to the size of single cells, which allows us to manipulate single cells and micrometer-sized particles with minimal disturbance in deep tissue. Ultrahigh frequency ultrasound based imaging improves spatial resolution to better identify abnormalities in tissue samples. We have developed intravascular ultrasound (IVUS) imaging transducers to improve resolution for detecting lesion on the lumen wall^12, 13^ and single cell intracellular delivery of various types of macromolecules into cell cytoplasm^5, 6^ using ultrahigh frequency transducers up to 150 MHz. Single cell acoustic trapping^14^ and cell signaling monitoring using transducers with the center frequency of up to 200 MHz was demonstrated ^3, 4, 14^.

The main and immediate questions using ultrahigh frequency ultrasonic transducers are the measurements of acoustic radiation force (ARF) at the focus of transducers. Current technology is not mature enough to measure ARF of ultrahigh frequency ultrasound with the center frequency of over 60 MHz ^15, 16^. Many applications using ultrahigh frequency ultrasound only refers to the input voltages and duty factor without measuring actual ARF at focus. Numerical simulation using commercial packages was used to estimate ARF as an alternative.

Since we engineered immune cells and developed the next generation intracellular delivery technique using ultrasonic transducers, our lab standardized transducer’s specification such as aperture diameter, center frequency (*f_c_*), *f_number_*, and housing sizes to systemically optimize the performance of transducers for these applications to investigate biological effects of high frequency ultrasonic transducers to cells^5, 6, 17–20^.

Here, for the first time, we quantitatively measured ARF of ultrahigh frequency transducers which provides a better understanding of the mechanism of high frequency ultrasound based single cell manipulation. We estimated ARF of a pushing ultrasonic transducer (1, PUT in Fig. 1A-1B) using a model-based approach by mapping measured displacements of a solid sphere in gelatin phantom with theoretical predictions. Theoretical model of the motion of a solid sphere in a viscoelastic medium was developed and experimentally validated in our lab^21–24^. In our previous study, we developed a theoretical model of a gas bubble under ARF and measured the mechanical properties of the crystalline lens and the vitreous humor of bovine and porcine eyes using a 3 MHz transducer for pushing and a 25 MHz transducer for tracking^25–28^. In this study, we used a 130 MHz PUT to displace a solid sphere in a gelatin phantom and a 45 MHz tracking ultrasonic transducer (TUT) to track the displacement of the sphere. If this approach is successfully established to measure the ARF of ultrahigh frequency ultrasonic transducers, this approach will become a gold standard to be widely used in ultrasonic community.

**Fig. 1.**
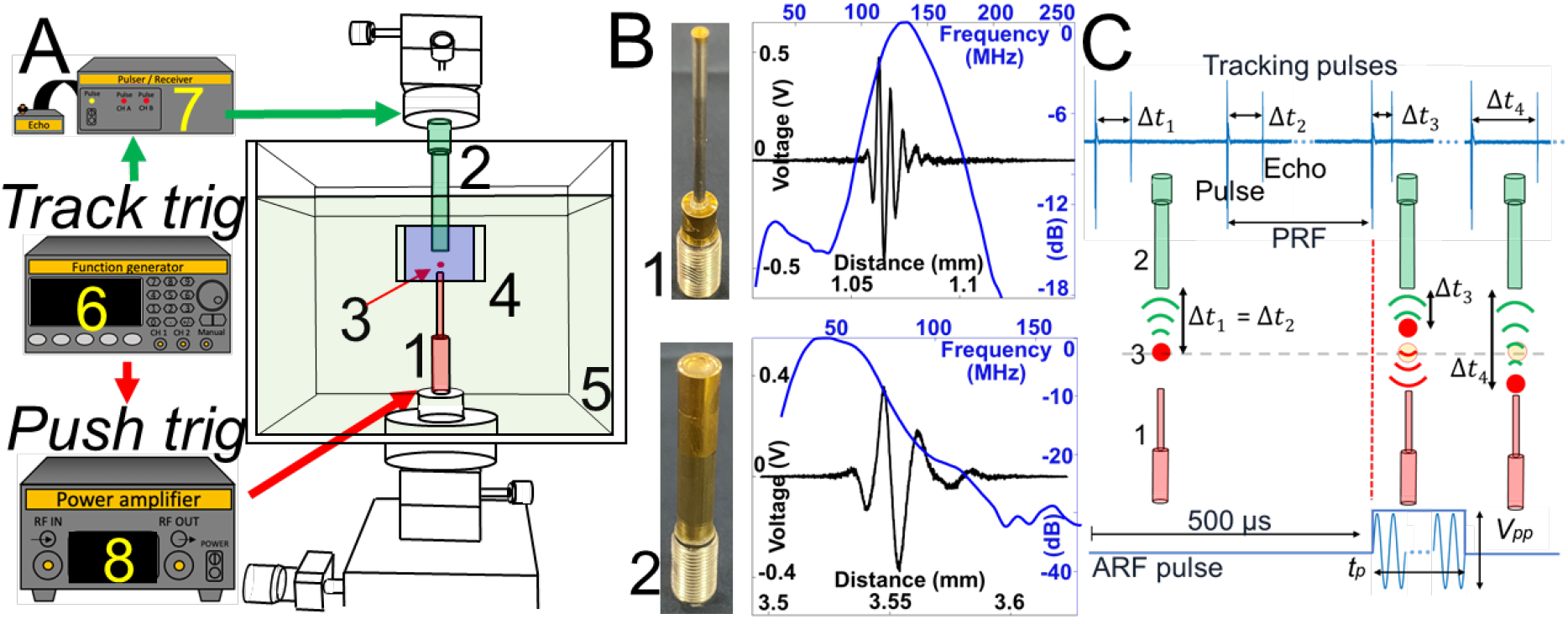
(A) Measurement system of acoustic radiation force (ARF) of ultrahigh frequency transducer is composed of 1. a pushing transducer (PUT) with (B) the center frequency of 130 MHz, 2. a tracking transducer (TUT) with (B) the center frequency of 45 MHz, 3. a solid sphere, 4. gelatin phantom block, 5. water cuvette, 6. function generator, 7. pulser / receiver, 8. power amplifier. Red arrows indicate a sequence for pushing triggering to generate ARF in (C) and green arrows indicate a sequence for tracking triggering to generate tracking pulses in (C). (C) When a sphere is moving under ARF from PUT, Δ*t* changes. *V_pp_* is peak-to-peak voltage and *t_p_* is pulse duration of ARF pulse. Pulse and echoes were saved for post-processing to reconstruct the displacement of a sphere ball using cross-correlation method.

To develop a system, we fabricated a PUT and TUT using lithium niobate plates (LNB, Boston Piezo-Optics) by following our lab’s protocol^12^. TEM was used to measure the thickness of LNB of PUT and it was approximately 10 μm, which matched our designed specification. Measured center frequency (*f_c_*) of PUT and IUT were 130 MHz and 45 MHz shown in Fig. 1B and Table I.

**TABLE 1.** Specification of pushing and imaging ultrasonic transducers.

| | $f_c$ (MHz) | LNB thickness ($\mu\text{m}$) | Aperture (mm) | $f_{number}$ | BW (%) |
| --- | --- | --- | --- | --- | --- |
| TUT | 45 | 50 | 3 | 1.16 | 80 |
| PUT | 150 | 10 | 1 | 1 | 58 |

We first placed a carbon steel sphere (CS) or a ruby sphere (Ruby) in a 2% gelatin phantom. We measured shear elastic modulus (*μ*) using Discovery Hybrid HR-20 rheometer (TA instruments). Measured shear elastic modulus of 2% gelatin phantom was 2300 ± 900 Pa from 10 specimens. PUT and TUT were carefully aligned using goniometer and translation stages with a CS or a Ruby sphere as shown in Fig. 1A. PUT was positioned in the gelatin phantom block with a sphere to minimize ARF loss due to reflection at the water and gelatin phantom boundary. After water cuvette was filled with water, PUT was connected to the power amplifier (8 in Fig. 1A) and TUT was connected to pulser / receiver (7 in Fig. 1A) and oscilloscope. A function generator (6 in Fig. 1A) controlled tracking and pushing triggering for radiofrequency (RF) data acquisition for post-processing to reconstruct the displacement of a sphere under ARF (Fig. 1C). Tracking pulses were emitted from TUT with 100 kHz pulse repetition frequency (PRF) and a pushing pulse was emitted from PUT with a pulse duration (*t_p_*) and peak-to-peak voltage (*V_pp_*) as shown in Fig. 2C. Five hundred pulse / echo were saved in a storage for post-processing to visualize the displacement of a sphere. Sampling frequency (*f_s_*) was 10 Gsamples/s and 50 million samples were saved to track the dynamics of the sphere for 5 ms. Cross correlation speckle tracking method was used to define displacement of the sphere^29^.

**Fig. 2.**
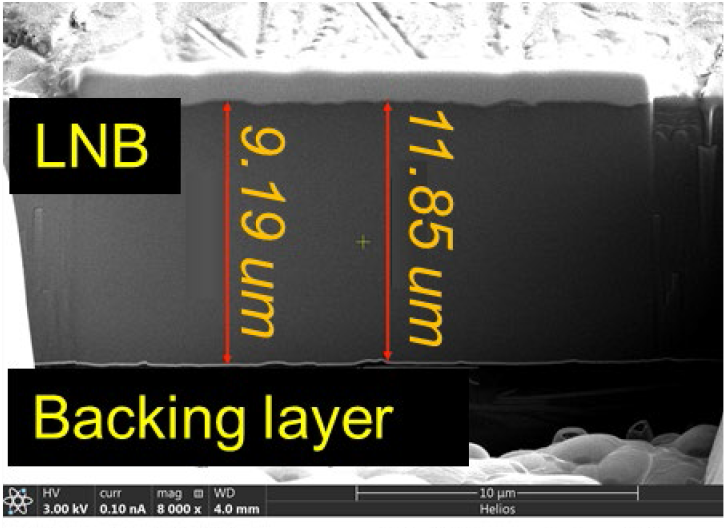
TEM image of LNB and backing layer of TUT

Previously developed and validated theoretical model formulates the dynamics of a microbubble in a viscoelastic medium under impulsive ARF. To use this theoretical model, we reasoned that the difference between gas bubbles and solid spheres is the density of material, which restricts breathing motion of microbubbles and makes the particle heavier. We changed the densities of air (1.2 kg/cm^3^) to densities of CS (7850 kg/cm^3^) and Ruby (4000 kg/cm^3^) to simulate solid sphere displacement. CS and Ruby models are limiting cases of microbubble theoretical model. Brief summary of a theory is the following. The governing equation of motion for an incompressible medium is

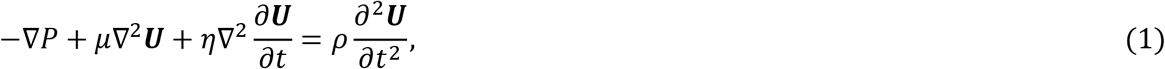

where *P* is internal pressure, ***U*** is the displacement vector of a sphere, *μ* and *η* are the shear elastic modulus and shear viscosity, respectively, *ρ* is medium density and *t* is time. The boundary conditions at the surface of a microbubble (*r* = *R*) are

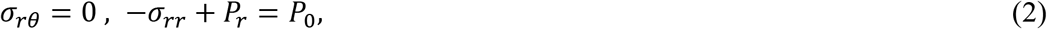

where *σ_rr_* and *σ_rθ_* are stress tensor components at the surface of the object, *R* is the radius of bubble or eventually the size of a solid sphere, *P_r_* is acoustic radiation pressure acting on the surface of the bubble and *P_0_* is an internal gas pressure^23^. Since pressure *P* and, consequently, *σ_rr_* are defined up to a constant, the internal gas pressure *P_0_* may be set equal to zero.

External force applied to a sphere (*F*^(*ext*)^) is a rectangular shape (Fig. 1C):

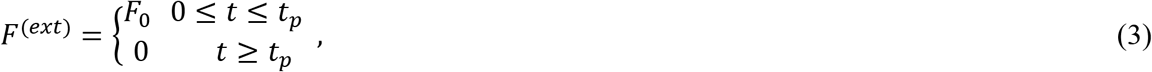

where *t_p_* is the duration of acoustic radiation pulse (Fig. 1C).

In this study, we changed *F*_0_ by changing *V_pp_* and *t_p_*. Now that we have the governing equation, boundary conditions, and external forces, Fourier transform of equations (1) – (3) was taken to find the displacement of a sphere under ARF in frequency domain. By taking inverse Fourier transform, time domain component of the displacement of a sphere in radial direction (*U_r_*) is

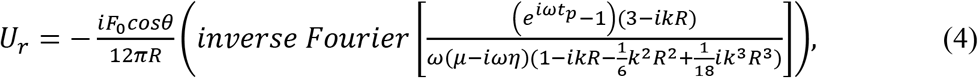

where *θ* is the angle between the bubble (solid sphere) movement direction and radial direction. Since we consider the displacement of a sphere along the ultrasound propagation, *θ* is set to zero. Therefore, *U_r_* is represents the displacement of a sphere under consideration.

We first used a 150 μm radius CS sphere to investigate the feasibility of mapping between measurements and theoretical calculations of the dynamics of a solid sphere under ARF generated by a 130 MHz transducer. To estimate *F*_0_ which will be converted into ARF, *t_p_* was fixed to 20 μs and *V_pp_* were increased from 30 mV to 90 mV. These values were chosen from our lab’s other publications^3, 5, 6^ for the intracellular delivery of macromolecules and acoustic single cell stimulation. Because the gain of the power amplifier is 50 dB, 30 mV and 90 mV represent 9 V to 28 V. Measured displacements of a CS sphere are plotted in Fig. 3A and theoretical calculation to match with the measured displacements using equation (4) is shown in Fig. 3B. After the matching between measurements and theory, *F*_0_ were estimated to be 60 μN/s, 40 μN/s, 24 μN/s, and 9 μN/s for 90 mV, 70 mV, 50 mV, and 30 mV, respectively. Because *F*_0_ in equation (4) is Newton per unit time, *t_p_* of 20 μs should be multiplied to estimated ARF. To confirm estimated *F*_0_, we fixed 70 mV and changed *t_p_* as 1 μs, 10 μs, 20 μs, and 30 μs. If *F*_0_ of 40 μN/s was correctly estimated, theory and measurement using different *t_p_* should match. As shown in Figs. 3C and 3D, measured displacements and theoretical calculations match for varying *t_p_* for *F*_0_ of 40 μN/s. Therefore, measured ARF of *V_pp_* of 90 mV, 70 mV, 50 mV, and 30 mV with fixed 20 μs are 1.2 nN, 0.8 nN, 0.48 nN, and 0.18 nN, respectively.

**Fig. 3.**
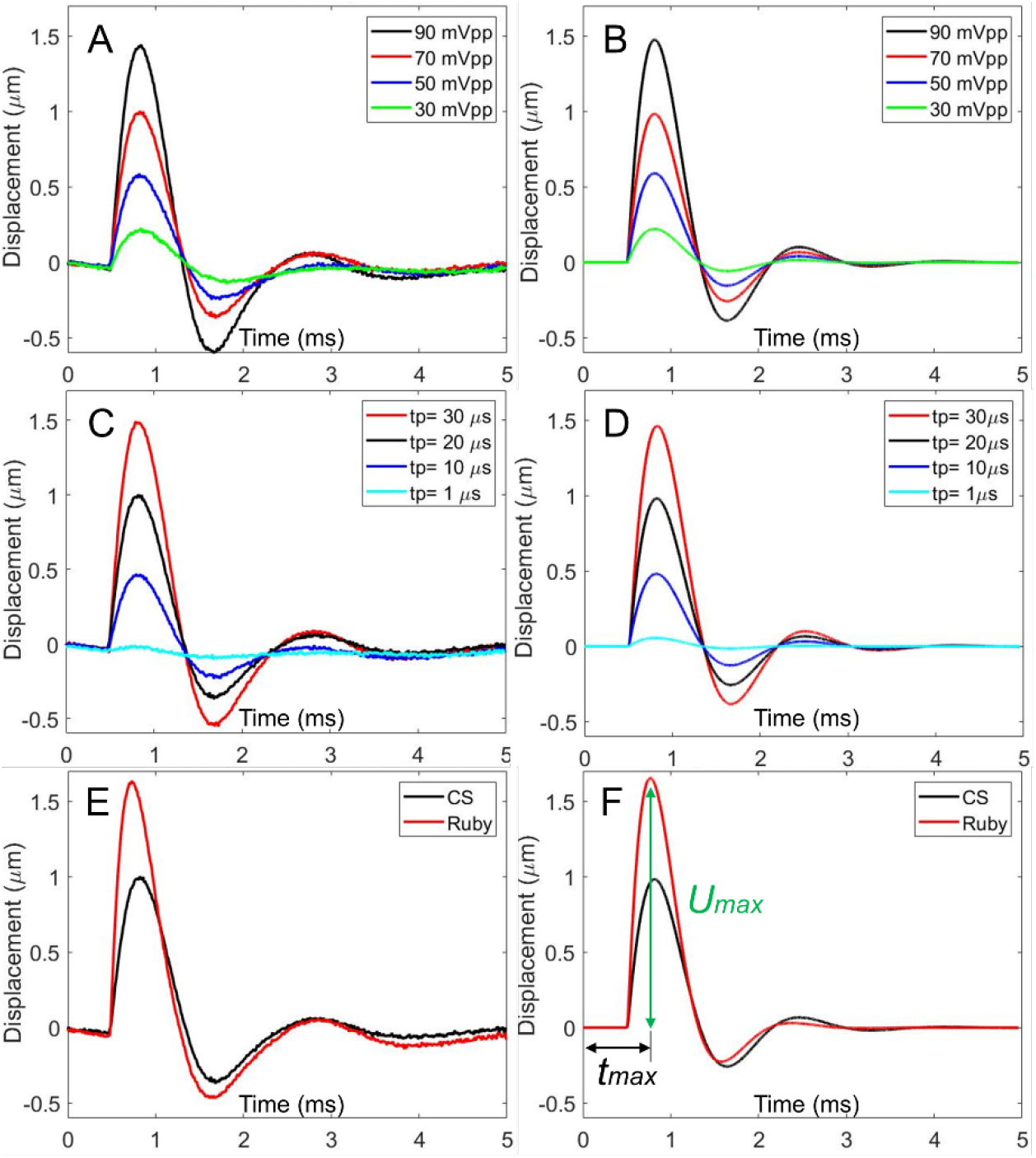
(A) Measured displacements and (B) theoretical calculations of a carbon steel sphere (CS) depending on *V_PP_*. Pulse duration (*t_p_*) is 20 μs. (C-D) To confirm mapping between measurements and a theory, *F*_0_ and *V_pp_* are fixed while *t_p_* is changed from 1 μs to 30 μs. (E-F) To confirm mapping between measurements and a theory, a ruby sphere (Ruby) and CS were compared. *U_max_* and *t_max_* represent maximum displacement of a sphere and time to reach *U_max_*.

As a second confirmation, we changed the material of a sphere. We used a 150 μm radius Ruby sphere and compared the displacements of a Ruby sphere with a CS sphere when *V_pp_* is 70 mV and *t_p_* is 20 μs. Considering the density of Ruby (4000 kg/cm^3^) and CS (7850 kg/cm^3^), time to reach maximum displacement (*t_max_*) was decreased and maximum displacement (*U_max_*) was increased as shown in Fig. 3E and 3F. Under the same ARF, lighter Ruby sphere travels longer distance within short time.

Because we confirmed this model-based ARF measurements by varying *V_pp_*, *t_p_*, and the material of a sphere, we measured ARF by comparing *U_max_* of a CS and a Ruby spheres within a design space of *V_pp_* of 30 mV, 50 mV, 70 mV, 90 mV and *t_p_* of 5 μs, 10 μs, 15 μs, 20 μs, 30 μs. We selected *V_pp_* and *t_p_*, which have been frequently used for intracellular delivery of macromolecules, acoustic tweezers, and single cell stimulations using ultrahigh frequency ultrasound. PUT, TUT, and a sphere were co-aligned in gelatin phantom not to cause any reflection and refraction which may occur when ultrasound waves are traveling across water and gelatin boundaries. Measured *F*_0_ are 9.9 ± 1.4 μN/s, 26.5 ± 4.2 μN/, 42.4 ± 3.0 μN/s, and 65.1 ± 5.8 μN/s for 30 mV, 50 mV, 70 mV, 90 mV, respectively. Therefore, estimated ARFs of PUT are ranging from 0.045 nN to 1.8 nM as shown in Fig. 4. We can conclude here that applied ARF to cells during intracellular delivery and single cell stimulation was in the range of several tens of pN to nN.

**Fig. 4.**
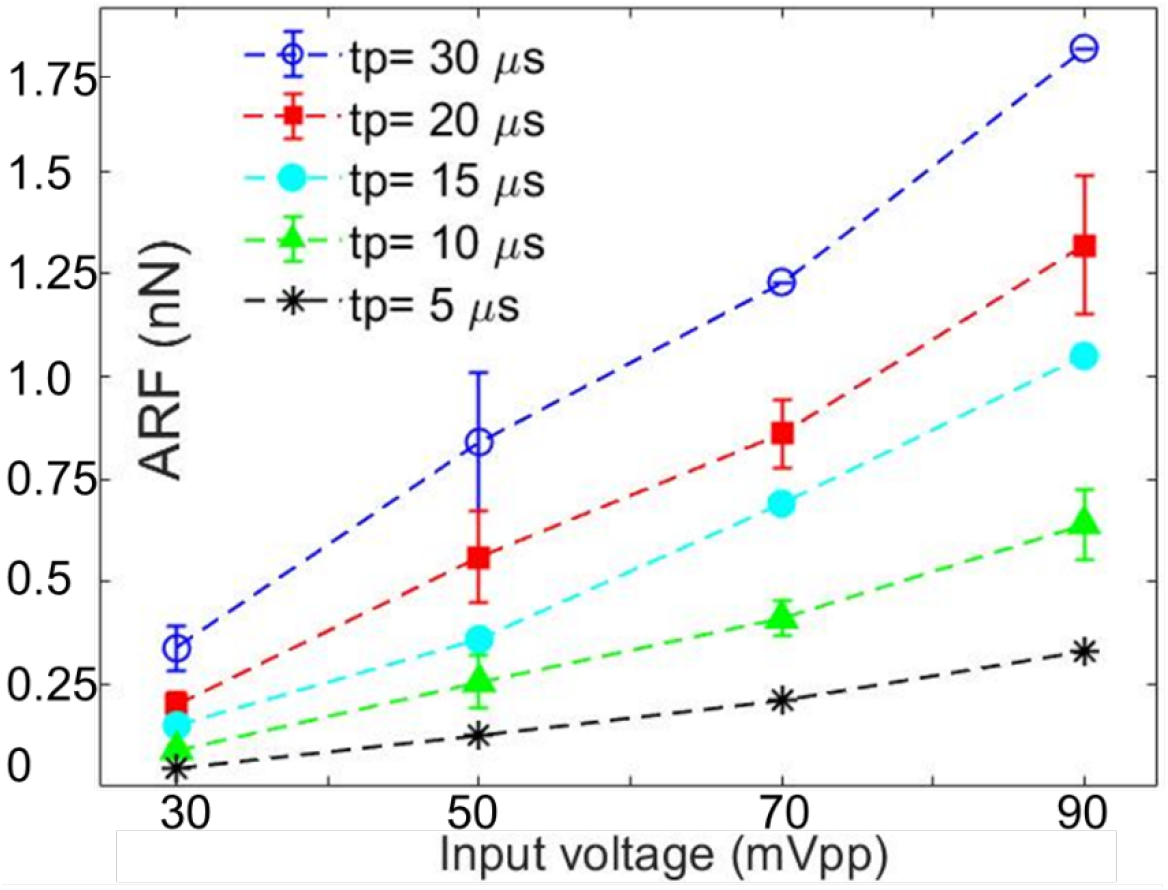
Measured ARF of 130 MHz ultrahigh frequency transducer is ranging from 0.045 nN to 1.8 nN

We used the PUT to demonstrate single cell stimulation and investigate calcium transport between cells. An integrated system was used to stimulate a target cell while capturing fluorescence resonance energy transfer (FRET) changes of the target cell and a neighboring cell to visualize calcium transport between two cells (Fig. 5A). The 3D stage was computer controlled with sub-micrometer precision, where the transducer was attached for focusing and stimulation of the target single cell. Foci of microscope and the transducer were coaligned for FRET based live cell imaging. To prevent standing waves between PUT aperture and the bottom of a cell culture dish, PUT was positioned at 45° with respect to cell culture dish (Fig. 5A). Particularly, standing waves induce pressure doubling at the rigid boundary condition.

**Fig. 5.**
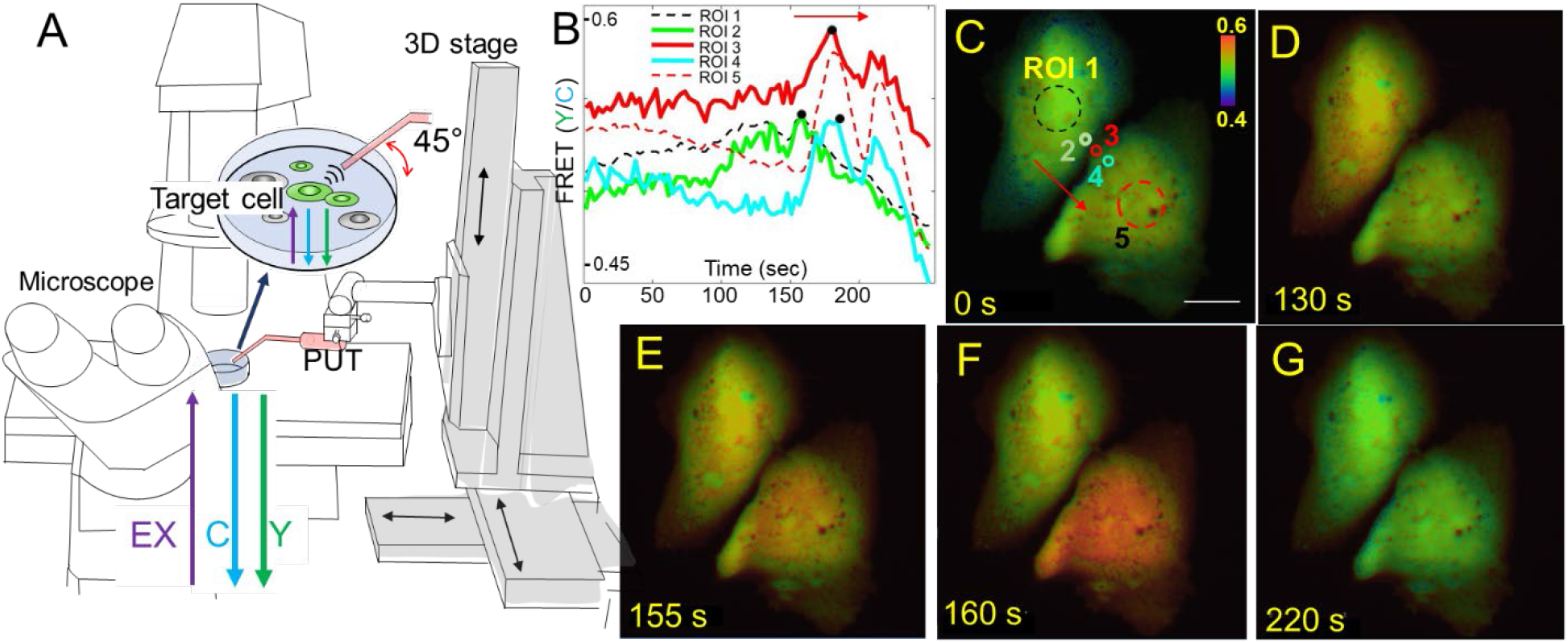
(A) PUT with 130 MHz center frequency stimulates a target cell expressing FRET biosensor to detect calcium transport between cells. PUT position is controlled by 3D stage. Microscope was used to image FRET changes of target cells after the stimulation using the intensity ratio between ECFP (C) and YPet (Y). (B-C) Time course changes of FRET of ROIs 1-5. ROI 1 is in the middle of the target cell and ROI 5 is in the middle of a neighboring cell. ROIs 2-4 are sequentially located from the stimulated target cell to the neighboring cell. Black dots indicate maximum FRET of ROIs 2-4. FRET changes of ROIs 2-4 indicate calcium transport as indicated red arrows and black dots on the FRET plot. A scale bar indicates 20 μm. The color scale bar in panel C represents the range of FRET ratio with purple and red colors, Indicating low and high levels of calcium concentration, respectively. (D-G) FRET images of two cells at the designated time.

Ultrasound pulses with *V_pp_* of 70 mV and *t_p_* of 5 μs were applied to the region of interest 1 (ROI 1) shown in Fig. 5C for 1 minute between 90 sec to 150 sec with a PRF of 1 kHz. From ARF measurements in Fig. 4, 0.21 nN ARF was applied every 1 ms for 1 minutes. FRET images were collected with the Ti2 Nikon fluorescence microscope and a cooled charge-coupled device (CCD) camera using the Nikon NIS-Elements AR software. Nikon NIS-Elements AR was used to calculate the pixel-by-pixel ratio images of FRET-YPet over ECFP after subtracting basal fluorescence level. In other words, FRET is the ratio between the intensity of YPet signal (Y in Fig. 5A) and ECFP signal (C in Fig. 5A), represented by a simple equation (FRET= Y/C) in Fig. 5A. High FRET value 0.6 in this case represents increased intracellular calcium concentration and low FRET value 0.4 indicates decreased and basal calcium level in cytoplasm of cells. Microscope filter setting for ECFP images was excitation (EX) 440/20 nm and emission (EM) 480/40 nm. FRET-YPet image filter setting was EX 440/20 nm and EM 535/30 nm. Calcium transport between two cells were clearly visualized due to time shift between maximum FRET in ROIs 2-3 (black dots in Fig. 5B and red arrows in Figs. 5B-C). Figs. 5C-G represent FRET images at 0 s, 130 s, 155 s, 160 s, and 220 s, respectively. First FRET increases only within the stimulated cell at Fig. 5D. Then, FRET increases in both cells in Fig. 5E. Quantitatively, the first FRET peak of ROI 1 is at 155 sec and the first FRET peak of ROI 5 is at 182 sec. Peak FRET of ROIs 2-4 shifts from 159 sec, 181 sec, and 184 sec, respectively. These results confirm calcium transport between two cells starting from the stimulated cell.

We developed an approach to measure ARF of ultrahigh frequency ultrasonic transducers using theoretical model of the dynamics of a sphere in a gelatin phantom. To estimate ARF at the focus of 130 MHz transducer, we matched measured *U_max_* with theoretical calculations. We selected appropriate ranges of *V_pp_* and *t_p_* for single cell applications and estimated ARFs were in the range of tens of pN to nN. FRET live cell imaging was performed to visualize calcium transport between cells after a target single cell was stimulated by the developed ultrasonic transducer. We conclude that calcium ion may be intracellularly delivered by acoustic radiation force in the range of pN to nN. Because the focal diameter of 130 MHz is approximately 10 μm, converted pressure needed for calcium ion intracellular delivery is 2.7 Pa. This study may establish a gold standard to estimate ARF of ultrahigh frequency transducers for single cell applications.

## Acknowledgments

This work was supported in part by National Institute of Health (NIH) grant No. GM120493 and National Science Foundation (NSF) grant No. CBET 1943852, 2019 AD&T Discovery Fund, and Harper Cancer Research Institute CCV grant.

## DATA AVAILABILITY

The data that support the findings of this study are available from the corresponding author upon reasonable request.

## References

1. Kim, M. G.; Park, J.; Lim, H. G.; Yoon, S.; Lee, C.; Chang, J. H.; Shung, K. K., Label-free analysis of the characteristics of a single cell trapped by acoustic tweezers. Sci Rep 2017, 7 (1), 14092.

2. Lee, C.; Lee, J.; Kim, H. H.; Teh, S. Y.; Lee, A.; Chung, I. Y.; Park, J. Y.; Shung, K. K., Microfluidic droplet sorting with a high frequency ultrasound beam. Lab Chip 2012, 12 (15), 2736–42.

3. Yoon, S.; Pan, Y.; Shung, K.; Wang, Y., FRET-Based Ca(2+) Biosensor Single Cell Imaging Interrogated by High-Frequency Ultrasound. Sensors (Basel) 2020, 20 (17).

4. Hwang, J. Y.; Yoon, C. W.; Lim, H. G.; Park, J. M.; Yoon, S.; Lee, J.; Shung, K. K., Acoustic tweezers for studying intracellular calcium signaling in SKBR-3 human breast cancer cells. Ultrasonics 2015, 63, 94–101.

5. Yoon, S.; Wang, P.; Peng, Q.; Wang, Y.; Shung, K. K., Acoustic-transfection for genomic manipulation of single-cells using high frequency ultrasound. Sci Rep 2017, 7 (1), 5275.

6. Yoon, S.; Kim, M. G.; Chiu, C. T.; Hwang, J. Y.; Kim, H. H.; Wang, Y.; Shung, K. K., Direct and sustained intracellular delivery of exogenous molecules using acoustic-transfection with high frequency ultrasound. Sci Rep 2016, 6, 20477.

7. Hwang, J. Y.; Lim, H. G.; Yoon, C. W.; Lam, K. H.; Yoon, S.; Lee, C.; Chiu, C. T.; Kang, B. J.; Kim, H. H.; Shung, K. K., Non-contact high-frequency ultrasound microbeam stimulation for studying mechanotransduction in human umbilical vein endothelial cells. Ultrasound Med Biol 2014, 40 (9), 2172–82.

8. Kalies, S.; Kuetemeyer, K.; Heisterkamp, A., Mechanisms of high-order photobleaching and its relationship to intracellular ablation. Biomed Opt Express 2011, 2 (4), 805–16.

9. Le Harzic, R.; Riemann, I.; König, K.; Wüllner, C.; Donitzky, C., Influence of femtosecond laser pulse irradiation on the viability of cells at 1035, 517, and 345nm. Journal of Applied Physics 2007, 102 (11), 114701.

10. Cannata, J. M.; Ritter, T. A.; Chen, W. H.; Silverman, R. H.; Shung, K. K., Design of efficient, broadband single-element (20-80 MHz) ultrasonic transducers for medical imaging applications. IEEE Trans Ultrason Ferroelectr Freq Control 2003, 50 (11), 1548–57.

11. Lam, K. H.; Ji, H. F.; Zheng, F.; Ren, W.; Zhou, Q.; Shung, K. K., Development of lead-free single-element ultrahigh frequency (170-320MHz) ultrasonic transducers. Ultrasonics 2013, 53 (5), 1033–8.

12. Yoon, S.; Kim, M. G.; Williams, J. A.; Yoon, C.; Kang, B. J.; Cabrera-Munoz, N.; Shung, K. K.; Kim, H. H., Dual-element needle transducer for intravascular ultrasound imaging. J Med Imaging (Bellingham) 2015, 2 (2), 027001.

13. Yoon, S.; Williams, J.; Kang, B. J.; Yoon, C.; Cabrera-Munoz, N.; Jeong, J. S.; Lee, S. G.; Shung, K. K.; Kim, H. H., Angled-focused 45 MHz PMN-PT single element transducer for intravascular ultrasound imaging. Sens Actuators A Phys 2015, 228, 16–22.

14. Lee, J.; Shung, K. K., Radiation forces exerted on arbitrarily located sphere by acoustic tweezer. J Acoust Soc Am 2006, 120 (2), 1084–94.

15. Nagle, S. M.; Sundar, G.; Schafer, M. E.; Harris, G. R.; Vaezy, S.; Gessert, J. M.; Howard, S. M.; Moore, M. K.; Eaton, R. M.; Food, U. S.; Drug, A., Challenges and regulatory considerations in the acoustic measurement of high-frequency (>20 MHz) ultrasound. J Ultrasound Med 2013, 32 (11), 1897–911.

16. Umchid, S.; Gopinath, R.; Srinivasan, K.; Lewin, P. A.; Daryoush, A. S.; Bansal, L.; El-Sherif, M., Development of calibration techniques for ultrasonic hydrophone probes in the frequency range from 1 to 100 MHz. Ultrasonics 2009, 49 (3), 306–11.

17. Pan, Y.; Yoon, S.; Zhu, L.; Wang, Y., Acoustic mechanogenetics. Current Opinion in Biomedical Engineering 2018, 7, 64–70.

18. Kim, M. G.; Yoon, S.; Kim, H. H.; Shung, K. K., Impedance matching network for high frequency ultrasonic transducer for cellular applications. Ultrasonics 2016, 65, 258–67.

19. Kim, M. G.; Yoon, S.; Chiu, C. T.; Shung, K. K., Investigation of Optimized Treatment Conditions for Acoustic-Transfection Technique for Intracellular Delivery of Macromolecules. Ultrasound Med Biol 2018, 44 (3), 622–634.

20. Pan, Y.; Yoon, S.; Sun, J.; Huang, Z.; Lee, C.; Allen, M.; Wu, Y.; Chang, Y. J.; Sadelain, M.; Shung, K. K.; Chien, S.; Wang, Y., Mechanogenetics for the remote and noninvasive control of cancer immunotherapy. Proc Natl Acad Sci U S A 2018, 115 (5), 992–997.

21. Karpiouk, A. B.; Aglyamov, S. R.; Ilinskii, Y. A.; Zabolotskaya, E. A.; Emelianov, S. Y., Assessment of shear modulus of tissue using ultrasound radiation force acting on a spherical acoustic inhomogeneity. IEEE Trans Ultrason Ferroelectr Freq Control 2009, 56 (11), 2380–7.

22. Aglyamov, S. R.; Karpiouk, A. B.; Ilinskii, Y. A.; Zabolotskaya, E. A.; Emelianov, S. Y., Motion of a solid sphere in a viscoelastic medium in response to applied acoustic radiation force: Theoretical analysis and experimental verification. J Acoust Soc Am 2007, 122 (4), 1927–36.

23. Ilinskii, Y. A.; Meegan, G. D.; Zabolotskaya, E. A.; Emelianov, S. Y., Gas bubble and solid sphere motion in elastic media in response to acoustic radiation force. J Acoust Soc Am 2005, 117 (4 Pt 1), 2338–46.

24. Aglyamov, S.; Karpiouk, A.; Mehrmohammadi, M.; Yoon, S.; Kim, S.; Ilinskii, Y.; Zabolotskaya, E.; Emelianov, S., Elasticity Imaging and Sensing Using Targeted Motion: From Macro to Nano. Current Medical Imaging 2012, 8 (1), 3–15.

25. Yoon, S.; Aglyamov, S.; Karpiouk, A.; Emelianov, S., Correspondence: Spatial variations of viscoelastic properties of porcine vitreous humors. IEEE Trans Ultrason Ferroelectr Freq Control 2013, 60 (11), 2453–60.

26. Yoon, S.; Aglyamov, S. R.; Karpiouk, A. B.; Kim, S.; Emelianov, S. Y., Estimation of mechanical properties of a viscoelastic medium using a laser-induced microbubble interrogated by an acoustic radiation force. J Acoust Soc Am 2011, 130 (4), 2241–8.

27. Yoon, S.; Aglyamov, S.; Karpiouk, A.; Emelianov, S., A high pulse repetition frequency ultrasound system for the ex vivo measurement of mechanical properties of crystalline lenses with laser-induced microbubbles interrogated by acoustic radiation force. Phys Med Biol 2012, 57 (15), 4871–84.

28. Yoon, S.; Aglyamov, S.; Karpiouk, A.; Emelianov, S., The mechanical properties of ex vivo bovine and porcine crystalline lenses: age-related changes and location-dependent variations. Ultrasound Med Biol 2013, 39 (6), 1120–7.

29. Lubinski, M. A.; Emelianov, S. Y.; O’Donnell, M., Speckle tracking methods for ultrasonic elasticity imaging using short-time correlation. IEEE Trans Ultrason Ferroelectr Freq Control 1999, 46 (1), 82–96.

